# Gut bacterial composition and predicted functional profiles associated with depressive symptom burden in Japanese adults recruited independently of depression diagnosis

**DOI:** 10.64898/2026.09.13.751222

**Authors:** Shunsuke Ichikawa, Aiko Hoshino, Shun Okuhara

## Abstract

Reported associations between depressive symptoms and the gut microbiota have been inconsistent, particularly with respect to predicted microbial function. We conducted an exploratory cross-sectional study of 17 adults to examine associations of depressive symptom burden with fecal bacterial composition and predicted functional potential. Depressive symptoms were assessed using the Japanese version of the Center for Epidemiologic Studies Depression Scale (CES-D); 13 participants scored <16 and 4 met the screening cutoff (≥16) for elevated depressive symptoms. Bacterial communities were profiled by 16S ribosomal RNA gene sequencing. Weighted UniFrac community structure differed between groups by permutational multivariate analysis of variance. Exploratory taxonomic analysis identified 31 features differentiating the groups. Among features enriched in the higher CES-D group, those assigned to *Veillonella*, *Parabacteroides*, and Porphyromonadaceae showed the highest linear discriminant analysis scores, whereas Firmicutes and several Lachnospiraceae- and Ruminococcaceae-related taxa were enriched in the lower CES-D group. PICRUSt2-predicted functional profiles differed between groups across four annotation systems. At the MetaCyc pathway level, the lower CES-D group showed nominally higher predicted purine salvage and several amino acid, phospholipid, and coenzyme A biosynthesis I pathways, whereas the higher CES-D group showed nominally higher predicted de novo pyrimidine deoxyribonucleotide biosynthesis, several TCA-cycle variants, glucose/xylose and N-acetylneuraminate degradation, the glycolysis/Entner-Doudoroff superpathway, and the urea cycle. These findings provide exploratory evidence that depressive symptom burden is associated with gut bacterial composition and predicted functional profiles among Japanese adults recruited independently of depression diagnosis.

## Introduction

Depressive disorders are a major contributor to non-fatal health loss worldwide, and their public-health importance is not confined to individuals who meet full diagnostic criteria ^1^. Population-based evidence indicates that subthreshold depressive symptoms are also associated with meaningful decrements in health, supporting assessment of depressive symptom burden as a continuum as well as by diagnostic or screening categories ^2^. The Center for Epidemiologic Studies Depression Scale (CES-D) was developed as a 20-item self-report instrument for epidemiologic research and measures the frequency of depressive symptoms during the preceding week; a validated Japanese version is available ^3,4^.

The microbiota–gut–brain axis provides a biologically plausible framework through which the intestinal microbial community could be associated with mood-related phenotypes.

Communication between the gut and the brain is bidirectional and may involve autonomic and enteric neural pathways, the vagus nerve, neuroendocrine signaling, the hypothalamic–pituitary–adrenal axis, immune and inflammatory processes, and intestinal and blood–brain barrier function ^5,6^. Microbial transformation of dietary and host-derived substrates can also generate short-chain fatty acids, tryptophan-derived metabolites, bile-acid derivatives, and other neuroactive or immunomodulatory compounds that may influence host physiology locally or systemically ^5,7^.

Experimental transfer studies add biological plausibility. Transplantation of fecal microbiota from patients with major depressive disorder into germ-free mice induced depressive-like behavioral changes and was accompanied by disturbances in microbial genes and host metabolites related to carbohydrate and amino acid metabolism ^8^. A separate transfer study similarly showed that fecal microbiota from patients with depression induced anhedonia- and anxiety-like behaviors and altered tryptophan metabolism when transferred to microbiota-depleted rats ^9^. Such findings demonstrate that a depression-associated microbial community can influence behavior and metabolism under controlled experimental conditions ^5,6^.

Human studies have increasingly linked depressive phenotypes to gut microbial community structure, although findings are heterogeneous. Systematic reviews and meta-analyses have reported inconsistent results for within-sample and between-sample diversity and limited agreement for individual taxa across studies ^10–12^. Nevertheless, recurrent signals include lower abundances of several taxa with the capacity to produce short-chain fatty acids, particularly *Faecalibacterium* and *Coprococcus*, and differences in members of Lachnospiraceae and Ruminococcaceae ^10–12^. In the Flemish Gut Flora Project, with validation in independent cohorts, *Faecalibacterium* and *Coprococcus* were consistently associated with better quality-of-life measures, while *Coprococcus* and *Dialister* were depleted among participants with depression ^13^. Two subsequent population-scale studies extended these observations to dimensional depressive symptom scores: one identified replicated associations with 13 taxa, including multiple Ruminococcaceae- and Lachnospiraceae-related lineages, and another found associations between microbial diversity, community composition, and depressive symptom levels across six ethnic groups ^14,15^.

At the same time, no single taxonomic signature has emerged as a reproducible marker of depression. Associations for broad phyla and common genera vary in direction, and even the most frequently reported taxa are not uniformly altered across cohorts ^11,12^. Differences in participant characteristics, geography and ethnicity, habitual diet, psychotropic and non-psychotropic medications, stool collection, sequencing region, taxonomic database, statistical model, and the definition of the depressive phenotype can all contribute to between-study variation ^12,13,15^. Large population-based analyses have likewise shown that medication use, diet, stool characteristics, and other host and environmental factors contribute to interindividual variation in gut microbiome composition ^16^.

Functional characterization may complement taxonomic profiling because taxonomically distinct microbial communities can encode overlapping core functions, while differing in pathway variants, substrate preferences, and the organisms contributing those functions ^17^. Population-scale metagenomic analysis has associated mental quality of life and depression with microbial modules related to γ-aminobutyric acid and the dopamine metabolite 3,4-dihydroxyphenylacetic acid, illustrating the potential value of examining microbial functional capacity rather than taxonomy alone ^13^. In patients with major depressive disorder, shotgun metagenomics combined with fecal metabolomics identified coordinated differences in bacterial species, microbial genes, and metabolites involving γ-aminobutyrate, phenylalanine, and tryptophan metabolism ^18^. More recent multi-omics work has reported concurrent alterations in microbial functions, circulating amino acids and bile acids, lipid metabolism, and immune cell profiles, further indicating that depression-associated signals may span several metabolic and host-response domains rather than a single pathway ^19^.

Compared with case–control studies of diagnosed major depressive disorder, studies that integrate taxonomic ecology and pathway-resolved functional analysis across the continuum of depressive symptoms remain less common ^11,14,15^. The present exploratory cross-sectional study therefore examined fecal bacterial community composition in adults assessed with the Japanese CES-D. We evaluated within-sample diversity, abundance-weighted phylogenetic community structure, and differentially represented taxa in participants below versus at or above the conventional CES-D cutoff of 16. We then used PICRUSt2 to compare predicted functional profiles summarized by COG, Enzyme Commission number, KEGG Orthology, and MetaCyc pathway annotations ^20^. Finally, to retain the information contained in the full range of symptom scores and to complement the cutoff-based comparison, we assessed associations between predicted MetaCyc pathway relative abundances and continuous CES-D scores ^21^. The objective was to identify coordinated taxonomic and predicted functional features associated with recent depressive symptom burden and to generate testable targets for confirmation in larger, well-controlled cohorts using direct functional and metabolomic measurements.

## Materials and Methods

### Ethics approval and informed consent

The Ethics Committee of Nagoya University Graduate School of Medicine approved the study protocol (approval No. 2024-0265). All procedures were conducted in accordance with the approved protocol and the principles of the Declaration of Helsinki. All participants provided written informed consent before participating in the study.

### Participants and stool collection

This exploratory cross-sectional study included 17 adults aged 21–46 years in Japan. Participants collected stool samples at home using a fecal sampling kit containing a preservation buffer for gut microbiota analysis (TechnoSuruga Laboratory, Shizuoka, Japan). According to the manufacturer, the buffer maintains fecal microbial community profiles for approximately 1 month at 1–30 °C, allowing samples to be transported at ambient temperature. Participants mailed the samples to the laboratory, where they were immediately frozen at −20 °C upon receipt and stored at that temperature until DNA extraction. The interval between stool collection and laboratory freezing was not recorded for individual participants; however, based on the collection and mailing procedure, it was expected to be approximately 2–3 days.

### Assessment of depressive symptoms

Depressive symptom burden during the preceding week was assessed using the validated Japanese version of the 20-item Center for Epidemiologic Studies Depression Scale (CES-D) ^3,4^. Each item was rated using four frequency categories: rarely or none of the time, 1–2 days, 3–4 days, or 5 or more days during the preceding week. The 16 negatively worded items were scored from 0 to 3 in ascending order of frequency, whereas the four positively worded items (items 4, 8, 12, and 16) were reverse-scored from 3 to 0. Item scores were summed to obtain a total score ranging from 0 to 60, with higher scores indicating a greater depressive symptom burden. The Japanese wording and scoring direction of each item are provided in Table S1. Participants were classified using the conventional CES-D cutoff of 16: those with total scores of 16 or higher were assigned to the higher depressive symptom group, whereas those with scores below 16 were assigned to the control group. This classification was used for analytical purposes and was not interpreted as a clinical diagnosis of depression.

### Microbiome profiling and bioinformatic analysis

Microbial DNA was extracted from stool samples using the NucleoSpin DNA Stool kit (Macherey-Nagel, Düren, Germany) with bead beating. The V3–V4 region of the bacterial 16S rRNA gene was amplified by PCR. The resulting amplicons were purified and indexed by a second PCR, and the completed libraries were quantified and assessed for quality before sequencing. Paired-end sequencing was performed on an Illumina MiSeq platform using the MiSeq Reagent Kit v3 to generate 2 × 300-bp reads.

Following primer removal, quality filtering, and paired-end read merging, the sequences were processed using QIIME 2 (v2023.7) ^22^. The DADA2 plugin was used to remove noise and chimeric sequences and to generate amplicon sequence variants (ASVs) ^23^. Taxonomy was assigned to representative ASV sequences using the QIIME 2 feature-classifier plugin with the EzBioCloud 16S database as the reference ^24^. Representative sequences were aligned and used to construct a phylogenetic tree.

Microbial diversity was analyzed using the QIIME 2 diversity plugin. Alpha diversity was evaluated using observed features, Chao1 richness, Shannon diversity, Pielou’s evenness, and Faith’s phylogenetic diversity. Beta diversity was assessed using weighted UniFrac distances ^25^, visualized by principal coordinate analysis, and compared between the control and higher depressive symptom groups using permutational multivariate analysis of variance. Differentially represented taxa between the groups were identified using linear discriminant analysis effect size (LEfSe; v1.0.8) ^26^.

Predicted microbial functional profiles were generated from the ASV abundance table and representative sequences using PICRUSt2 (v2.4.1). Functional profiles were summarized according to Enzyme Commission numbers, KEGG Orthology, Clusters of Orthologous Groups, and MetaCyc pathways. Differences in predicted functional composition between the groups were evaluated using Bray–Curtis distances and permutational multivariate analysis of variance, with principal coordinate analysis used for visualization. Group-level differences in individual MetaCyc pathways were examined using STAMP (v2.1.3) ^27^. Associations between MetaCyc pathway abundances and continuous CES-D scores were evaluated separately. Detailed laboratory procedures and bioinformatic parameter settings are provided in the Supplementary Methods.

## Results

### Participant characteristics and CES-D score distribution

Participant characteristics and CES-D-based group classification are summarized in Table 1, with detailed self-reported antibiotic-use information and participant-level CES-D item scores provided in Tables S2 and S3, respectively. The study included 17 participants aged 21–46 years, with a median age of 31 years. Eight participants were female and nine were male. Antibiotic use before fecal sample receipt was reported by three participants, whereas relevant antibiotic-use information was not reported by the remaining participants. CES-D total scores ranged from 0 to 26, with a median score of 11. Based on the prespecified cutoff score of 16, thirteen participants were classified as the control group and four were classified as the higher depressive symptom group (Table 1).

**Table 1.**
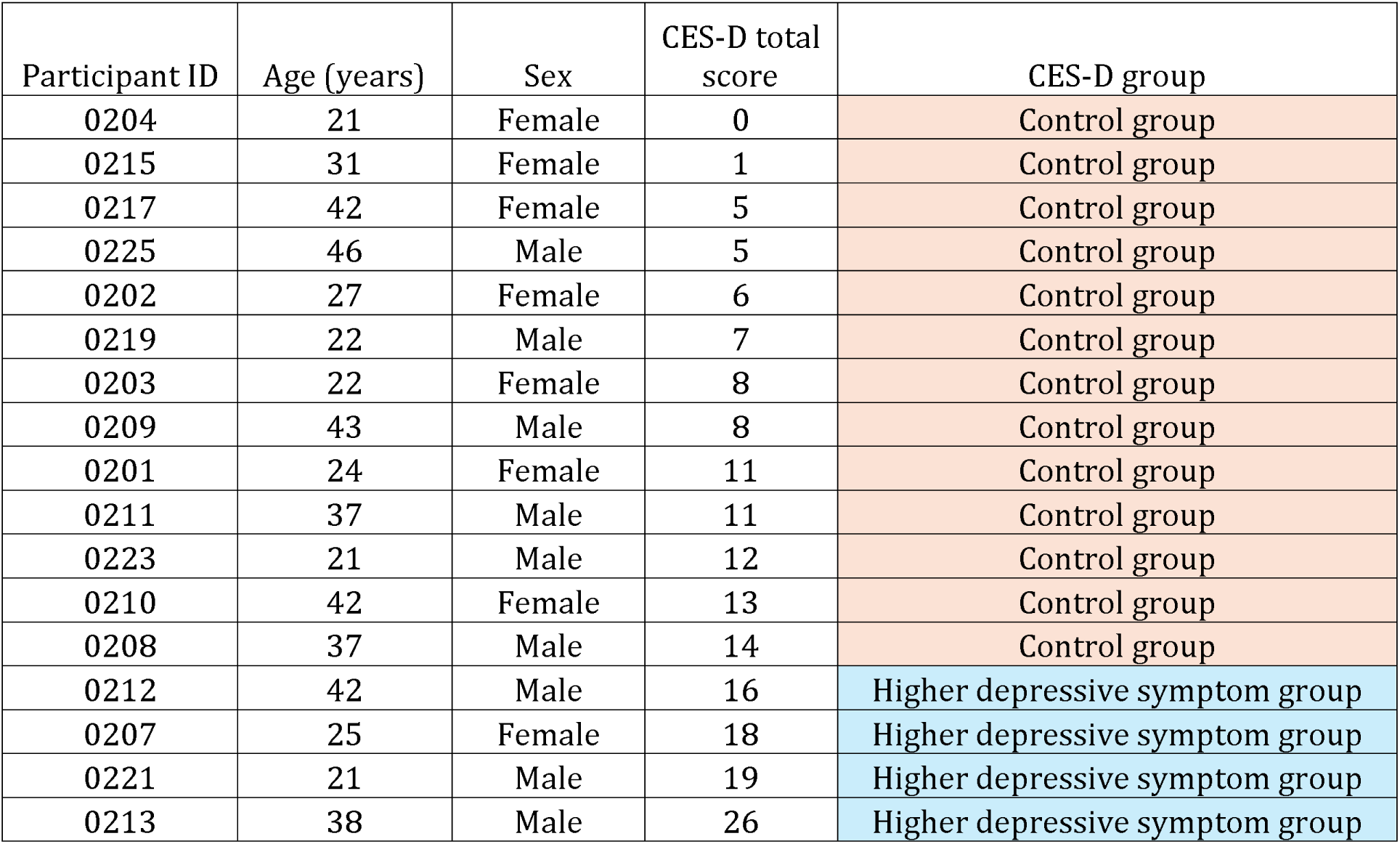
Participant characteristics and CES-D-based group classification.

We next examined whether CES-D scores were associated with age, sex, or self-reported antibiotic use. CES-D total scores showed no significant correlation with age (Spearman’s ρ = −0.019, p = 0.944). Age also did not differ between the control group and the higher depressive symptom group, with median ages of 31.0 and 31.5 years, respectively (Mann–Whitney U test, p = 0.955). Sex was not significantly associated with CES-D group assignment (Fisher’s exact test, p = 0.576). When CES-D total scores were analyzed as a continuous variable, the median score was 7.0 in female participants and 12.0 in male participants, and this difference was not statistically significant (Mann–Whitney U test, p = 0.123). Self-reported antibiotic use was also evaluated as a potential participant-level factor. All three participants with reported antibiotic use were classified into the control group, whereas none of the participants in the higher depressive symptom group reported antibiotic use. This distribution was not statistically significant (Fisher’s exact test, p = 0.541).

### Descriptive taxonomic profiles of the gut microbiome showed group-level differences between the control and higher depressive symptom groups

After sequence processing, the ASV table contained 1,341 ASVs across 17 fecal samples (Table S4). The total number of reads per sample ranged from 29,002 to 47,998, with a median of 38,103 reads. The number of observed ASVs per sample ranged from 93 to 372, with a median of 150. In the control group, the number of observed ASVs ranged from 105 to 372, with a median of 154, whereas the higher depressive symptom group showed 93 to 158 observed ASVs, with a median of 149.

Taxonomic composition of the gut microbiome was descriptively compared between the control group and the higher depressive symptom group at the phylum, order, and genus levels (Fig. 1). At the phylum level, Firmicutes and Bacteroidetes were the predominant phyla in both groups. The control group showed a higher mean relative abundance of Firmicutes than the higher depressive symptom group (57.8% vs. 39.7%), whereas the higher depressive symptom group showed a higher mean relative abundance of Bacteroidetes (41.3% vs. 31.6%). Actinobacteria also showed a higher mean relative abundance in the higher depressive symptom group than in the control group (11.4% vs. 6.1%). Proteobacteria showed comparable mean relative abundances between the control group and the higher depressive symptom group (3.5% vs. 3.9%). Verrucomicrobia was low in the control group but was detected at a higher mean relative abundance in the higher depressive symptom group (0.05% vs. 3.1%).

**Fig. 1.**
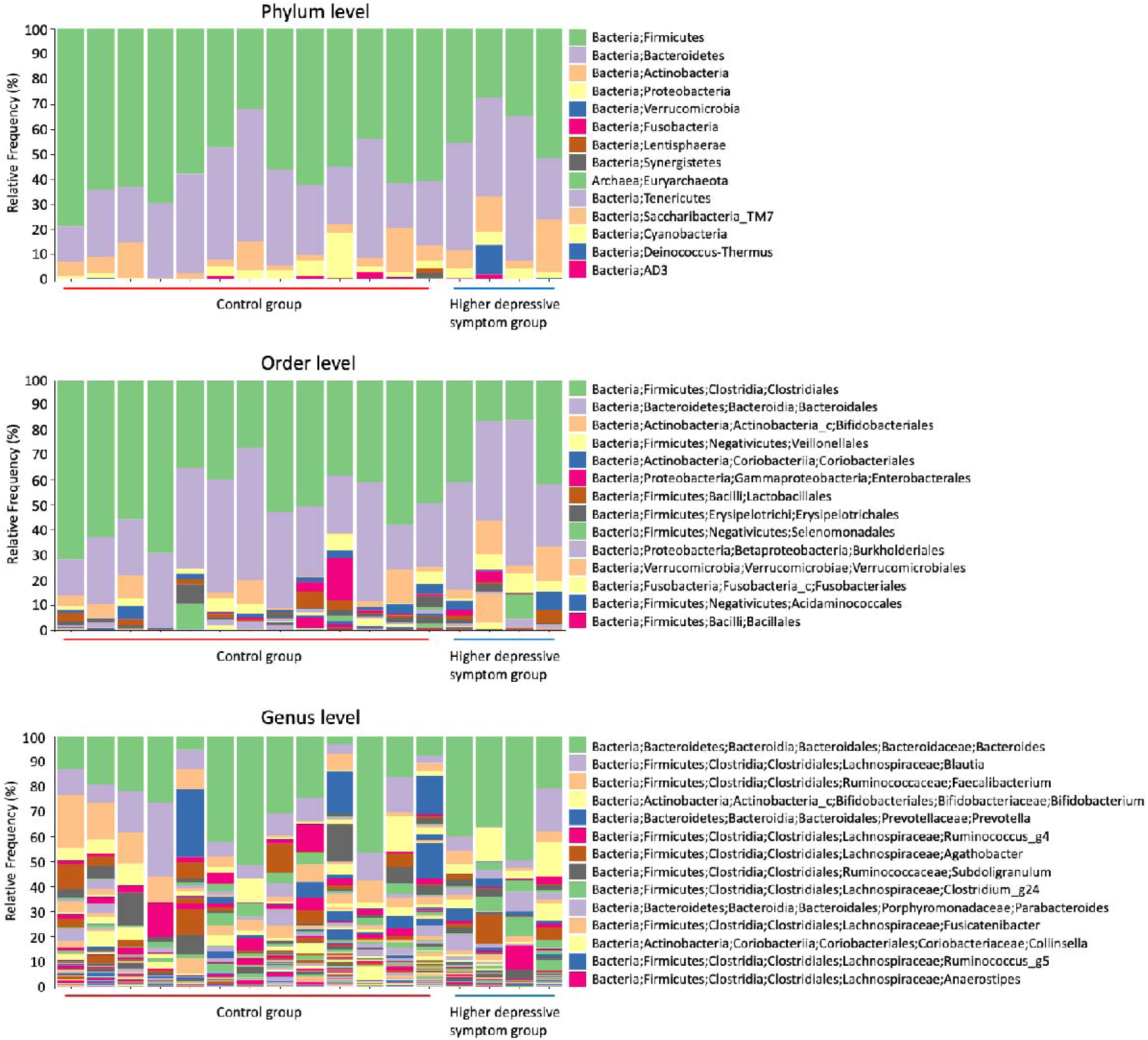
Taxonomic composition of gut microbiomes in the control and higher depressive symptom groups

At the order level, Clostridiales and Bacteroidales were the dominant orders across the cohort. Clostridiales accounted for a lower mean relative abundance in the higher depressive symptom group than in the control group (28.8% vs. 50.1%), whereas Bacteroidales accounted for a higher mean relative abundance in the higher depressive symptom group (41.3% vs. 31.6%). The higher depressive symptom group also showed higher mean relative abundances of Bifidobacteriales (8.6% vs. 3.9%), Veillonellales (4.8% vs. 2.1%), and Verrucomicrobiales (3.1% vs. 0.04%). In contrast, Erysipelotrichales and Acidaminococcales were slightly lower in the higher depressive symptom group than in the control group.

At the genus level, *Bacteroides* was the most abundant genus in both groups and showed a higher mean relative abundance in the higher depressive symptom group than in the control group (36.6% vs. 23.6%). *Bifidobacterium*, *Parabacteroides*, *Veillonella*, and *Akkermansia* also showed higher mean relative abundances in the higher depressive symptom group. In contrast, several genera within Firmicutes were lower in the higher depressive symptom group, including *Blautia* (6.5% vs. 10.3%), *Faecalibacterium* (2.6% vs. 7.1%), *Ruminococcus*_g4 (0.9% vs. 3.3%), *Agathobacter* (0.8% vs. 3.0%), *Subdoligranulum* (1.2% vs. 2.8%), and *Fusicatenibacter* (0.8% vs. 2.2%). *Prevotella* was also lower in the higher depressive symptom group than in the control group (0.07% vs. 4.6%).

These comparisons describe group-level mean relative abundances and do not indicate statistical significance; taxa identified as differentially abundant by LEfSe are presented separately below.

### Alpha and beta diversity analyses revealed altered gut microbial community structure without significant differences in within-sample diversity

Alpha diversity was compared between the control group and the higher depressive symptom group using five indices: observed features, Chao1 richness, Shannon diversity, Pielou’s evenness, and Faith’s phylogenetic diversity (Fig. S1). Rarefied alpha-diversity estimates were summarized at 26,666 reads per sample, which retained all 17 samples. The higher depressive symptom group showed numerically lower values across all five alpha-diversity indices than the control group; however, none of these differences reached statistical significance (observed features, p = 0.0799; Chao1 richness, p = 0.0822; Shannon diversity, p = 0.1194; Pielou’s evenness, p = 0.1963; Faith’s phylogenetic diversity, p = 0.1826). These results indicate that no statistically significant group difference in within-sample microbial diversity was detected in this cohort.

Beta diversity was then assessed using weighted UniFrac distances (Fig. S2). PERMANOVA detected a significant difference in abundance-weighted phylogenetic community composition between the control group and the higher depressive symptom group (pseudo-F = 2.42, p = 0.040, q = 0.040). These results suggest that the higher depressive symptom group differed from the control group in overall gut microbial community structure, despite the absence of statistically significant differences in alpha diversity.

### LEfSe analysis identified taxa associated with higher depressive symptoms

Differentially abundant taxa between the control group and the higher depressive symptom group were further examined using linear discriminant analysis effect size (LEfSe) analysis ^26^ (Fig. 2 and Table S5). LEfSe identified 31 discriminative taxonomic features across multiple taxonomic levels. Of these, 22 features were enriched in the higher depressive symptom group, whereas 9 features were enriched in the control group.

**Fig. 2.**
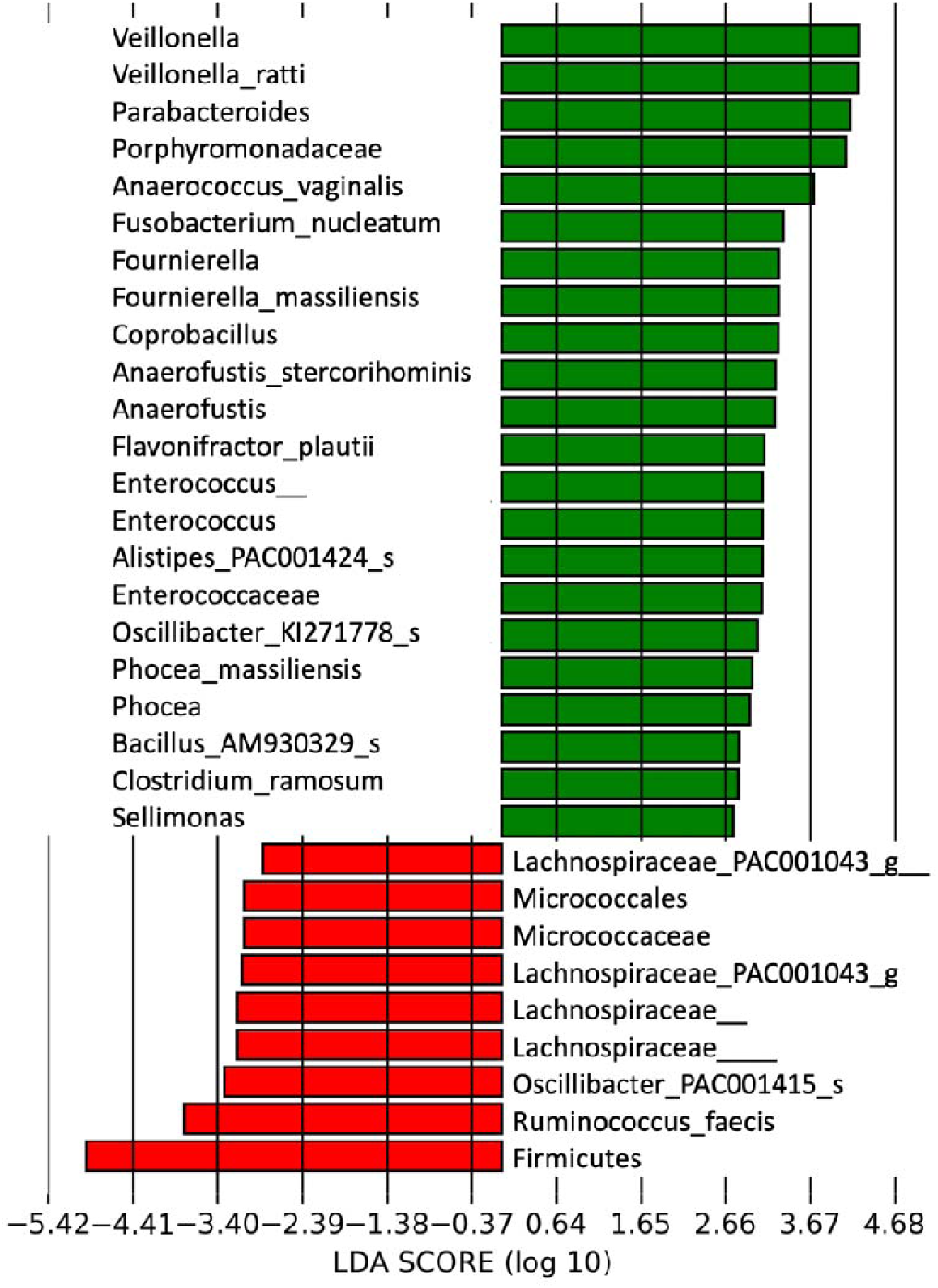
Differentially abundant gut microbial taxa identified by LEfSe analysis Green bars indicate taxa enriched in the higher depressive symptom group, whereas red bars indicate taxa enriched in the control group

In the higher depressive symptom group, the features with the highest LDA scores included *Veillonella*, *Veillonella ratti*, *Parabacteroides*, and Porphyromonadaceae. Additional taxa enriched in the higher depressive symptom group included *Anaerococcus vaginalis*, *Fusobacterium nucleatum*, *Fournierella*, *Fournierella massiliensis*, *Coprobacillus*, *Anaerofustis*, *Anaerofustis stercorihominis*, *Flavonifractor plautii*, *Enterococcus*, Enterococcaceae, *Phocea*, *Phocea massiliensis*, *Clostridium ramosum*, and *Sellimonas*.

In contrast, the control group was characterized by enrichment of Firmicutes and several Firmicutes-associated features, including *Ruminococcus faecis*, *Oscillibacter* PAC001415_s, and Lachnospiraceae-related taxa. Micrococcales and Micrococcaceae were also enriched in the control group.

Collectively, the taxa identified by LEfSe across both groups represented several bacterial lineages, including Firmicutes, Bacteroidetes, Fusobacteria, and Actinobacteria. The LEfSe cladogram further illustrated these group-associated lineages, highlighting Porphyromonadaceae and Enterococcaceae in the higher depressive symptom group and Micrococcales and Micrococcaceae in the control group (Fig. S3).

Overall, the LEfSe analysis indicated that the higher depressive symptom group was characterized by enrichment of selected Bacteroidetes- and Firmicutes-associated taxa, including *Veillonella*, *Parabacteroides*, and *Enterococcus*-related lineages, whereas the control group showed enrichment of Firmicutes and Lachnospiraceae/Ruminococcaceae-associated taxa. These findings are consistent with the beta-diversity result, which suggested group-level differences in abundance-weighted phylogenetic community composition.

### PICRUSt2-inferred microbial functional profiles differed between groups and showed pathway-level metabolic shifts

PICRUSt2 was used to infer microbial functional profiles based on EC number, KEGG Orthology, COG, and MetaCyc pathway annotations ^20^ (Table S6-S9). Principal coordinate analysis was then used to visualize differences in predicted functional profiles between the control group and the higher depressive symptom group. PERMANOVA based on Bray–Curtis distances detected significant group differences across all four functional annotation levels: COG (pseudo-F = 3.26, R² = 0.179, p = 0.0218), EC number (pseudo-F = 3.05, R² = 0.169, p = 0.0244), KEGG Orthology (pseudo-F = 3.07, R² = 0.170, p = 0.0210), and MetaCyc pathway (pseudo-F = 3.05, R² = 0.169, p = 0.0185) (Fig. S4). These results suggest that PICRUSt2-inferred microbial functional profiles differed between the control group and the higher depressive symptom group.

The MetaCyc pathway output included predicted abundance profiles for 389 pathways across the 17 fecal samples (Table S9), and these pathways were further compared between the control group and the higher depressive symptom group (Fig. 3 and Table S10). Among the predicted pathways, the Calvin-Benson-Bassham cycle showed a significantly higher mean relative abundance in the control group than in the higher depressive symptom group after multiple-testing correction (control, 0.879%; higher depressive symptom group, 0.795%; mean difference, 0.0836 percentage points; corrected p = 0.0315). Among the 35 pathways showing nominal between-group differences before multiple-testing correction, those with higher predicted relative abundances in the control group included adenine and adenosine salvage III, the non-oxidative branch of the pentose phosphate pathway, glycolysis III, several amino acid and phospholipid biosynthesis pathways, coenzyme A biosynthesis I, and tRNA charging. In contrast, those with higher predicted relative abundances in the higher depressive symptom group included de novo pyrimidine deoxyribonucleotide biosynthesis pathways, several variants of the tricarboxylic acid cycle, the superpathway of glucose and xylose degradation, the superpathway of glycolysis and Entner–Doudoroff, and the urea cycle (Fig. 3 and Table S10).

**Fig. 3.**
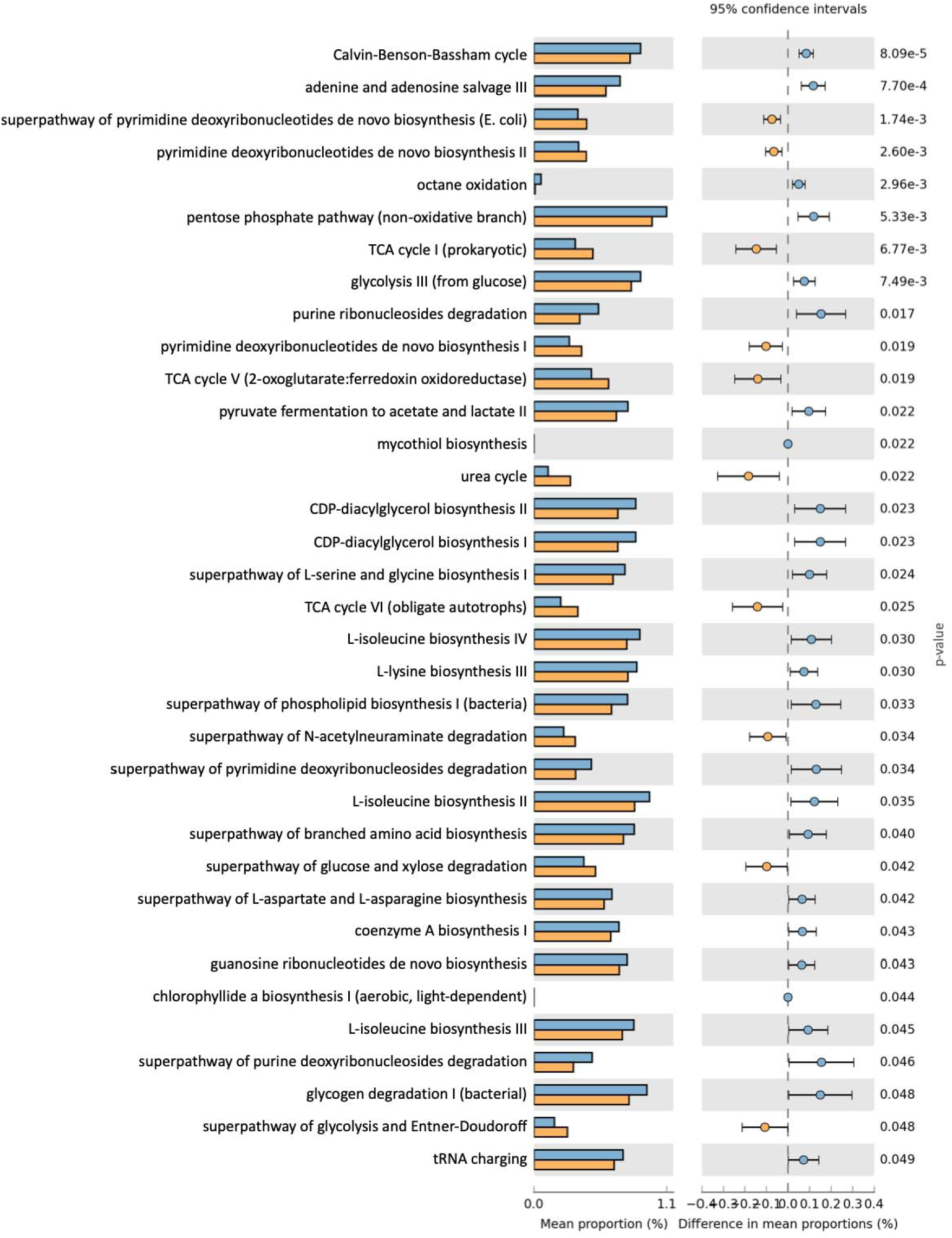
Differentially represented PICRUSt2-inferred MetaCyc pathways between the control and higher depressive symptom groups Orange dots indicate functions enriched in the higher depressive symptom group, whereas blue dots indicate functions enriched in the control group

Because dichotomizing CES-D scores at the 16-point cutoff produced a markedly unbalanced comparison of 13 participants with lower scores and four participants with higher scores and discarded variation within each group, we additionally analyzed CES-D total score as a continuous measure of depressive symptom burden. Spearman correlation analysis identified nominal associations with 31 of the 389 predicted MetaCyc pathways (*P* < 0.05). However, none remained significant after Benjamini–Hochberg correction across all 389 pathways (*q* < 0.05). The strongest association was observed for PWY-6969, TCA cycle V (2-oxoglutarate:ferredoxin oxidoreductase), which was positively correlated with CES-D score (ρ = 0.749, permutation *P* = 0.000830, *q* = 0.143). Restricting the analysis to the 354 pathways detected in at least five participants did not alter the multiple-testing conclusion.

Comparison of the continuous-score and binary-group analyses showed partial concordance between the two approaches. Of the 31 pathways with nominal associations with continuous CES-D score, seven overlapped with the 35 pathways previously highlighted in the binary-group comparison. These comprised PWY-6969, TCA cycle V; PWY0-166, the superpathway of pyrimidine deoxyribonucleotides de novo biosynthesis; PWY-7184, pyrimidine deoxyribonucleotides de novo biosynthesis I; TCA, TCA cycle I; PWY-7187, pyrimidine deoxyribonucleotides de novo biosynthesis II; PWY-6609, adenine and adenosine salvage III; and CALVIN-PWY, the Calvin–Benson–Bassham cycle. For all seven pathways, the direction of the continuous association was consistent with the corresponding difference between the control and higher depressive symptom groups.

Predicted functional profiles differed between the lower and higher CES-D groups at both the Kyoto Encyclopedia of Genes and Genomes ortholog (KO) and Enzyme Commission (EC) levels. PERMDISP detected no differences in multivariate dispersion between the groups at either the KO or EC level (P = 0.552 and P = 0.709, respectively). Continuous analyses also showed associations between CES-D scores and both KO (pseudo-F = 3.204, R² = 0.176, P = 0.023) and EC profiles (pseudo-F = 2.928, R² = 0.163, P = 0.036).

At the individual-feature level, 472 of 6,836 KOs and 175 of 2,047 ECs were nominally associated with CES-D scores, whereas 402 KOs and 147 ECs showed nominal differences between the groups (Table S7 and S8). None remained significant after Benjamini–Hochberg correction. The relative abundance of superpathway of N-acetylneuraminate degradation (P441-PWY) was higher in the higher depressive symptom group than in the control group (Fig. 3 and Table S10). Among ECs selected as candidates for mucin-glycan deconstruction, sialate O-acetylesterase (EC 3.1.1.53; ρ = 0.507, *P* = 0.039) and α-N-acetylglucosaminidase (EC 3.2.1.50; ρ = 0.486, *P* = 0.049) showed nominal positive associations with CES-D scores ^28^. Comparable nominal associations were observed for the corresponding KOs, K05970 (ρ = 0.516, *P* = 0.036) and K01205 (ρ = 0.513, *P* = 0.037) ^29^.

In addition, the combined relative abundance of β-galactosidase and β-N-acetylhexosaminidase, which remove terminal residues from diverse glycans, was 1.170% in the higher depressive symptom group and 0.873% in the control group. This combined measure was positively associated with CES-D scores (ρ = 0.505, permutation *P* = 0.041). In contrast, exo-α-sialidase, α-L-fucosidase, arylsulfatase, endo-α-N-acetylgalactosaminidase, and β-N-acetylhexosaminidase were not individually associated with CES-D scores at *P* < 0.05. These findings did not demonstrate an overall increase in mucin-degradation capacity. Nevertheless, the exploratory results suggested that some predicted enzymatic functions involved in the terminal processing of glycans, including mucin O-glycans, may increase as CES-D scores rise.

## Discussion

16S rRNA gene sequencing remains a practical approach for exploratory studies of community composition. PICRUSt2 uses phylogenetic placement of marker-gene sequences and annotated reference genomes to infer gene-family content and reconstruct metabolic pathways, including MetaCyc pathways ^20^. This approach can therefore be used to examine whether taxonomic differences are accompanied by differences in predicted functional potential. Taken together, the nominal pathway differences suggest a broad shift in the predicted functional repertoire of the bacterial community rather than a uniform alteration in a single depression-related pathway.

Previous studies support the general relevance of nucleotide metabolism to depression but do not provide a consistent directional model. Altered circulating inosine, hypoxanthine, xanthine, urate, and other purine metabolites have been reported in patients with major depressive disorder ^30^, whereas a plasma metabolomic study of older adults suggested downregulated purine metabolism in depression ^31^. An antidepressant-naïve 16S study also identified altered predicted pathways involving purine degradation and amino acid metabolism in depressive patients ^32^. In addition, combined serum and urine metabolomics identified disturbances in host purine and pyrimidine metabolism in unmedicated patients with major depressive disorder ^33^. These studies establish nucleotide metabolism as a potentially relevant metabolic domain, but circulating or urinary host metabolites cannot be equated with bacterial nucleotide pathways predicted from 16S data. Thus, the present findings do not determine whether bacterial nucleotide metabolism alters host purine or pyrimidine availability.

The nominal enrichment of the non-oxidative pentose phosphate pathway in the control group may be related to the generation and interconversion of sugar-phosphate precursors. This reversible branch connects pentose metabolism with glycolysis and can provide ribose-5-phosphate for nucleotide synthesis and other biosynthetic intermediates; importantly, unlike the oxidative branch, it does not itself generate NADPH or metabolic energy. A previous human 16S study identified the pentose phosphate pathway and starch and sucrose metabolism as microbiome functions associated with major depressive disorder, although the authors emphasized the need for biochemical and metabolomic confirmation ^34^. In adolescents with depression, inferred glycolysis/gluconeogenesis functions were also reported to differ from controls ^35^. These observations provide partial external support for an association between depressive phenotypes and microbial central carbon metabolism, but they do not establish a conserved direction of change.

The predicted relative abundances of several carbohydrate-degradation pathways, including the superpathway of glucose and xylose degradation, were nominally higher in the higher depressive symptom group. This finding may be ecologically related to the higher abundance of *Bacteroides* and related taxa observed in this group, because some *Bacteroides* species encode dedicated polysaccharide-utilization loci that enable the metabolism of complex dietary glycans ^36^. Human shotgun metagenomic and fecal metabolomic analysis has similarly identified increased *Bacteroides* and decreased *Blautia* and *Eubacterium* in major depressive disorder, together with disturbances involving carbohydrate, nucleotide, lipid, and amino acid metabolites ^18^. Another shotgun study found increased *Bacteroides* and decreased *Faecalibacterium*, *Ruminococcus*, and *Eubacterium* in moderate or severe depression, with functional differences involving carbohydrate- and host-glycan-utilization genes ^37^. Furthermore, transplantation of microbiota from patients with major depressive disorder into microbiota-depleted mice induced depression-like behavior accompanied by disturbances in carbohydrate and amino acid metabolism ^8^.

The nominally higher abundance of glycolysis III in controls, together with higher TCA-cycle variants in the higher depressive symptom group, does not indicate a simple shift from glycolysis to oxidative energy production. MetaCyc includes several taxon-specific, partial, oxidative, reductive, and alternative TCA-cycle configurations that can operate under different environmental conditions. Human metabolomic studies have reported increased circulating pyruvate and decreased citrate in major depressive disorder, indicating altered host energy metabolism ^38^. More recent shotgun metagenomic and targeted metabolomic work also found disturbances in host glycolysis, TCA-cycle, and ornithine-cycle metabolites associated with depressive severity ^39^. These findings support the broad relevance of energy metabolism to depression but cannot be used to infer that the bacterial TCA variants detected here reflect mitochondrial dysfunction or increased energy production in the participants.

Amino acid metabolism provides an important point of contact with previous depression research. Several predicted amino acid biosynthesis pathways were nominally higher in the control group. Human shotgun metagenomic and fecal metabolomic analyses have identified coordinated changes in microbial genes and metabolites related to GABA, phenylalanine, and tryptophan metabolism in major depressive disorder ^18^. Independent shotgun sequencing has also demonstrated taxonomic and tryptophan-pathway differences between patients with major depressive disorder and controls ^40^. Population-scale analyses have associated depressive symptoms or mental quality of life with the microbial potential for GABA, glutamate, serotonin-related, and dopamine-metabolite production ^13,14^.

The nominally higher phospholipid and CDP-diacylglycerol biosynthesis pathways in the control group may likewise indicate greater predicted capacity for bacterial membrane biogenesis. Depression-related multi-omics studies have reported disturbances in host serum glycerophospholipid metabolism ^33^, and fecal microbiota transplantation from women with prenatal depression to germ-free mice has been associated with depressive-like behavior and altered host glycerophospholipid and sphingolipid networks ^41^. These findings make lipid metabolism biologically relevant to the microbiota–gut–brain axis.

The nominally higher predicted relative abundance of the urea cycle in the higher depressive symptom group may reflect differences in microbial nitrogen handling. A human metagenomic and serum metabolomic study found enrichment of the urease accessory protein UreE in participants with depression, with *Enterobacter cloacae* proposed as an important contributor ^42^. Disturbances in host ornithine-cycle metabolites have also been associated with depressive severity ^39^.

The taxonomic findings provide additional ecological context. The control group showed higher relative abundances of several members of *Faecalibacterium*, *Blautia*, *Ruminococcus*, *Agathobacter*, and *Subdoligranulum*. Population studies and systematic reviews have frequently, although not universally, reported depletion of *Faecalibacterium*, *Coprococcus*, *Subdoligranulum*, and related Ruminococcaceae or Lachnospiraceae taxa in association with depression or higher depressive symptom scores ^10,12–14^. A large shotgun metagenomic study further demonstrated depletion of *Faecalibacterium prausnitzii* and pathways related to short-chain fatty acid production among adults with depressive symptoms ^43^. This provides partial ecological consistency with the direction observed in the present cohort.

Finally, PICRUSt2 predicts gene-family and pathway potential from marker-gene sequences and annotated reference genomes; it does not directly measure microbial genes, transcription, protein abundance, metabolite concentrations, or pathway flux ^20^. The cross-sectional design also precludes determining whether microbial differences contributed to depressive symptoms or resulted from symptom-associated differences in diet, sleep, medication, intestinal physiology, or other behaviors. The small and unequal group sizes, and the fact that reported antibiotic exposure occurred only among controls, represent additional potential sources of instability and confounding, because antibiotic perturbation can rapidly reduce microbial diversity and alter community composition, with recovery sometimes remaining incomplete ^44^. Overall, the results support an exploratory association between CES-D-assessed depressive symptom burden and the predicted functional composition of the gut bacterial community. The control-associated profile emphasized nucleotide recycling and general biosynthetic functions, whereas the higher depressive symptom-associated profile emphasized de novo pyrimidine nucleotide synthesis and alternative central carbon, carbohydrate, and nitrogen metabolism. Confirmation will require a larger cohort, adjustment for relevant confounders, taxon-contribution analysis, shotgun metagenomics, and direct fecal or circulating metabolomic measurements.

## Statements and Declarations

### Ethics approval

The Ethics Committee of Nagoya University Graduate School of Medicine approved the study protocol (approval No. 2024-0265).

### Consent to participate

Written informed consent was obtained from all participants.

### Consent to publish

Not applicable. This manuscript does not contain any individual person’s identifiable data.

### Data availability

The raw 16S rRNA gene sequencing dataset and associated metadata have been deposited in the NCBI Sequence Read Archive (SRA) under BioProject accession number PRJNA1524636. Processed feature tables, taxonomic assignments, and analysis scripts are available from the corresponding author upon reasonable request.

## Abbreviations

ASV: amplicon sequence variant
CES-D: Center for Epidemiologic Studies Depression Scale
COG: Clusters of Orthologous Groups
EC: Enzyme Commission
GABA: γ-aminobutyric acid
KEGG: Kyoto Encyclopedia of Genes and Genomes
KO: KEGG Orthology
LDA: linear discriminant analysis
LEfSe: linear discriminant analysis effect size
PCoA: principal coordinate analysis
PERMANOVA: permutational multivariate analysis of variance
PICRUSt2: Phylogenetic Investigation of Communities by Reconstruction of Unobserved States 2

## Acknowledgements

We sincerely thank all study participants for providing fecal samples and completing the CES-D questionnaire.

## Author contributions

S.I. and A.H. conceptualized and designed the study, collected the data, and drafted and revised the manuscript. S.I. and S.O. analyzed and interpreted the data. All authors reviewed and approved the final manuscript for submission.

---

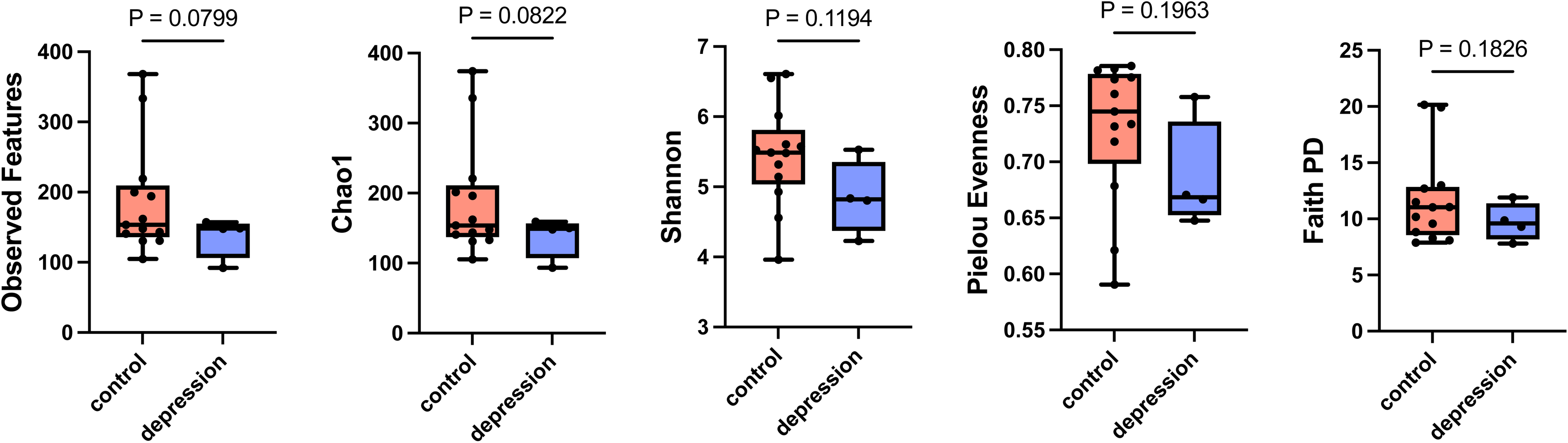

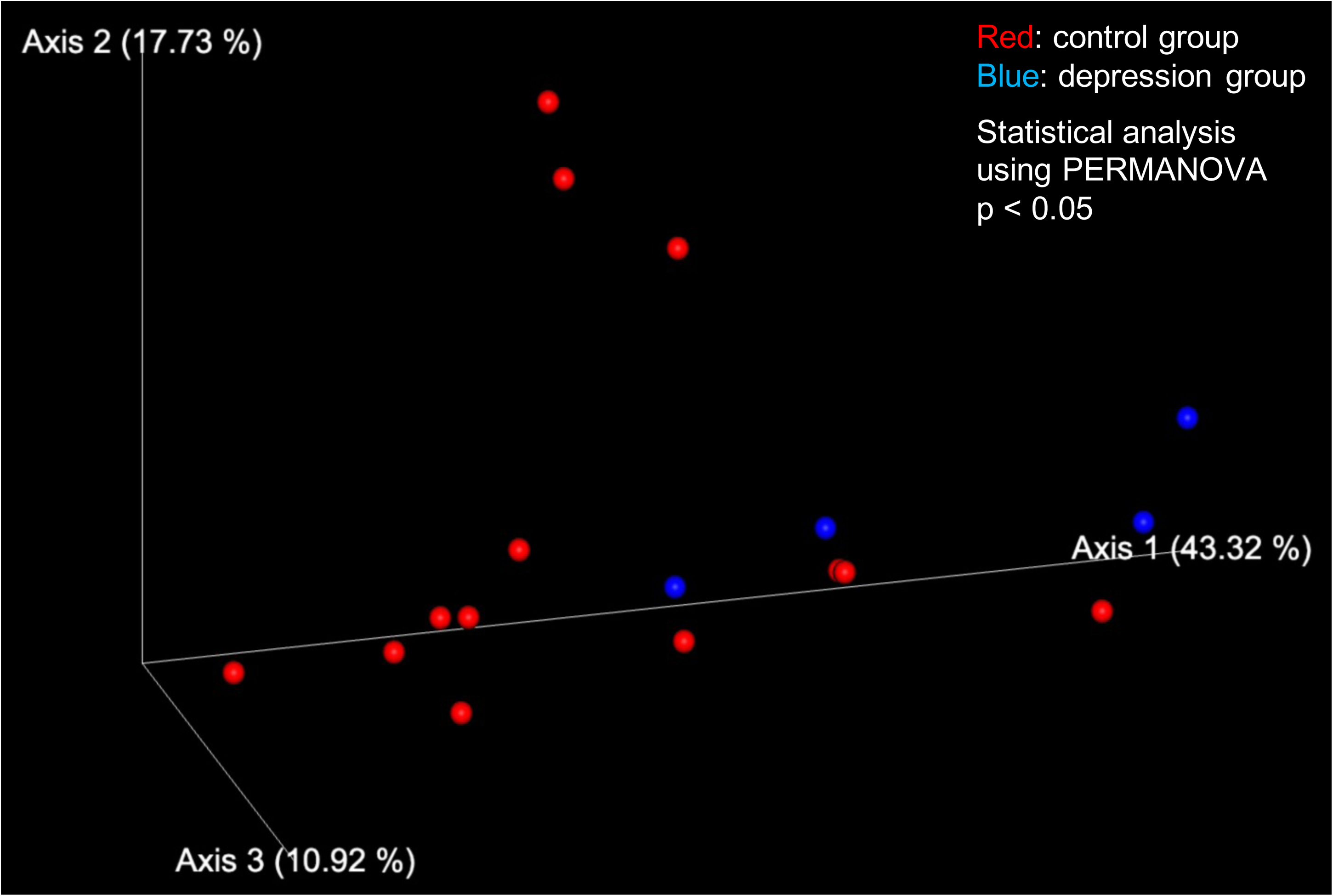

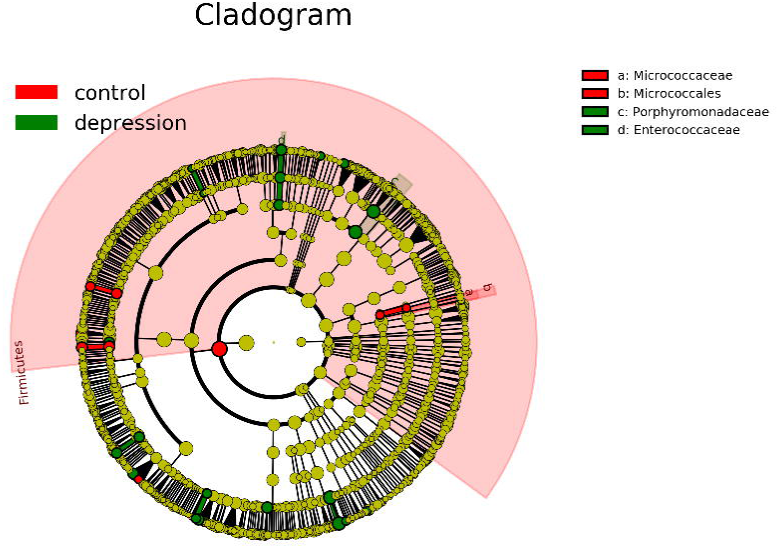

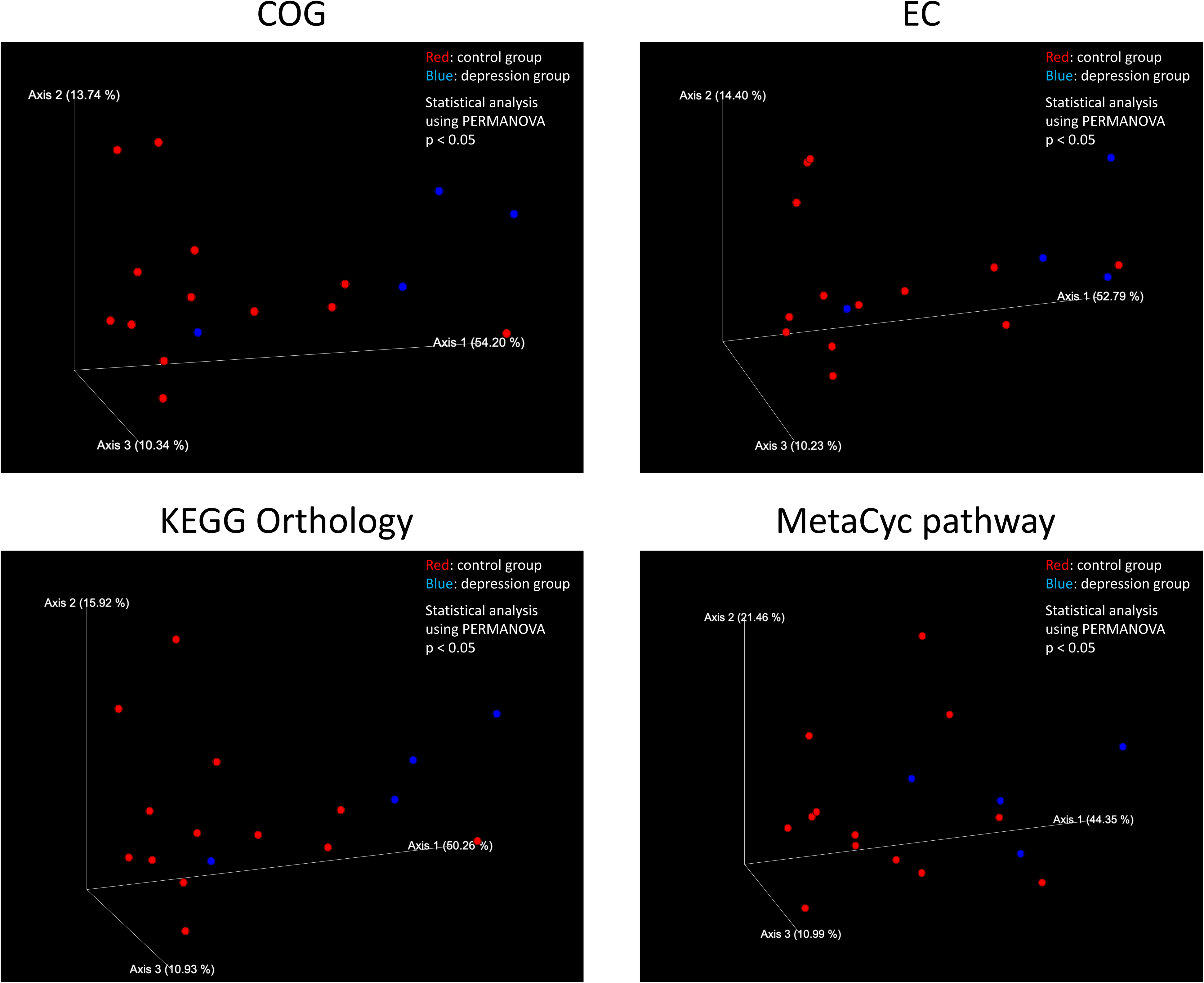

