## Supplementary information for "Gut bacterial composition and predicted functional profiles associated with depressive symptom burden in Japanese adults recruited independently of depression diagnosis"

**Supplementary Methods**

### DNA extraction from stool samples

An aliquot (220 µL) of each fecal suspension was transferred to an MN Bead Tube type A (Macherey-Nagel, Düren, Germany), and 850 µL of Buffer ST1 from the NucleoSpin DNA Stool kit (Macherey-Nagel) was added. The tube was mixed horizontally for 2–3 s to resuspend the sample and incubated at 70 °C for 5 min. Mechanical disruption was then performed at room temperature using a ShakeMan 6 homogenizer (Bio Medical Science, Tokyo, Japan) for six 90-s cycles at 2,500 rpm, with 30-s intervals between cycles.

After homogenization, the tube was centrifuged at 13,000 × g for 3 min. A 600-µL aliquot of the supernatant was transferred to a new 2-mL tube, and total DNA was subsequently purified using the NucleoSpin DNA Stool kit according to the manufacturer’s protocol.

### 16S rRNA gene amplification, library preparation, and sequencing

The V3–V4 region of the bacterial 16S rRNA gene was amplified from the extracted stool DNA using KOD One PCR Master Mix (Toyobo, Osaka, Japan) and the primer sets shown in Table SM1. Each primer comprised an Illumina adapter sequence, an N-mix spacer segment, and a V3–V4-specific sequence. Thermal cycling consisted of 25 cycles of denaturation at 98 °C for 10 s, annealing at 55 °C for 5 s, and extension at 68 °C for 15 s.

First-round PCR products were purified by adding VAHTS DNA Clean Beads (Vazyme) at 1.0 times the PCR reaction volume. Purified products were quantified using a Synergy H1 microplate reader (Agilent Technologies) and the QuantiFluor dsDNA System (Promega). Indexed sequencing libraries were then generated by a second PCR using KOD FX Neo polymerase (Toyobo) (Table SM2).

Table SM1. Primer sequences used for first-round amplification

| **Primer** | **Sequence (5′–3′)** |
| --- | --- |
| Forward | ACACTCTTTCCCTACACGACGCTCTTCCGATCT-NNNNN-CCTACGGGNGGCWGCAG |
| Reverse | GTGACTGGAGTTCAGACGTGTGCTCTTCCGATCT-NNNNN-GACTACHVGGGTATCTAATCC |

Table SM2. Composition of the second-round PCR mixture

| **Component** | **Volume per reaction** |
| --- | --- |
| 2× PCR Buffer for KOD FX Neo | 5.0 µL |
| dNTPs (2 mM each) | 2.0 µL |
| Forward primer (5 µM) | 0.5 µL |
| Reverse primer (5 µM) | 0.5 µL |
| Purified first-round PCR product | 1.0 µL |
| KOD FX Neo (1.0 U/µL) | 0.2 µL |
| Nuclease-free water | 0.8 µL |
| Total reaction volume | 10.0 µL |

The second-round PCR cycling profile comprised an initial denaturation at 94 °C for 2 min; repeated denaturation, annealing, and extension steps at 98 °C for 10 s, 60 °C for 30 s, and 68 °C for 30 s, respectively; and a final extension at 68 °C for 2 min. Indexed products were purified with VAHTS DNA Clean Beads at 1.0 times the PCR reaction volume. Library concentrations were measured using the Synergy H1 and QuantiFluor dsDNA System, and library quality was evaluated using a Fragment Analyzer with the dsDNA 915 Reagent Kit (Agilent Technologies). Paired-end sequencing was performed on an Illumina MiSeq system with the MiSeq Reagent Kit v3 to generate 2 × 300-bp reads.

### Sequence preprocessing and taxonomic assignment

Reads beginning with an exact match to the corresponding primer sequence were selected using fastx_barcode_splitter in FASTX-Toolkit (v0.0.14). This procedure was repeated for all 36 combinations of the six forward and six reverse N-mix variants. Primer sequences were removed with fastx_trimmer. Quality trimming was then performed with Sickle (v1.33) using a quality threshold of Q20. Reads that were 130 bp or shorter after trimming, together with their paired reads, were discarded.

Paired-end reads were merged using FLASH (v1.2.11). For the V3–V4 amplicons, merging used a 280-bp read length and a 10-bp overlap for an expected merged length of 410 bp. The merged sequences were processed in QIIME 2 (v2023.7). The DADA2 plugin was used to remove chimeric and noisy sequences and to generate an amplicon sequence variant (ASV) abundance table and representative sequences.

Taxonomic classification was performed with the QIIME 2 feature-classifier plugin by comparison with the EzBioCloud 16S database. Representative sequences were aligned, and a phylogenetic tree was constructed using the QIIME 2 alignment and phylogeny plugins.

### Microbial diversity and differential taxonomic analysis

Alpha- and beta-diversity analyses were conducted using the QIIME 2 diversity plugin. Alpha diversity was summarized using observed features, Chao1 richness, Shannon diversity, Pielou’s evenness, and Faith’s phylogenetic diversity. Abundance-weighted differences in community composition were evaluated using weighted UniFrac distances. Principal coordinate analysis was used for visualization, and group differences between the control group and the higher depressive symptom group were assessed by permutational multivariate analysis of variance. Principal coordinate plots were generated using the Emperor plugin.

Taxonomic profiles were summarized at the phylum, order, and genus levels. Differentially represented lineages between the two CES-D-defined groups were identified using linear discriminant analysis effect size (LEfSe; v1.0.8).

### Prediction and comparison of microbial functional profiles

The ASV abundance table and representative sequences exported from QIIME 2 were used as inputs for PICRUSt2 (v2.4.1). Predicted functional profiles were summarized according to Enzyme Commission numbers, KEGG Orthology identifiers, Clusters of Orthologous Groups, and MetaCyc pathways. Bray–Curtis distances were calculated for each functional annotation table. Predicted functional composition was visualized by principal coordinate analysis and compared between the two CES-D-defined groups by permutational multivariate analysis of variance. Group-level comparisons of individual MetaCyc pathways were performed using STAMP (v2.1.3).

### Association analysis using continuous CES-D scores

To complement the cutoff-based comparison, associations between continuous CES-D total scores and PICRUSt2-predicted MetaCyc pathway profiles were examined across all 17 participants. For each sample, the predicted abundance of each of the 389 pathways was divided by the sum of the predicted abundances of all pathways in that sample. Spearman’s rank correlation coefficient was then calculated between each pathway’s relative abundance and the CES-D total score, with average ranks assigned to tied observations. No covariates were included in these exploratory analyses.

Two-sided Monte Carlo permutation tests were performed by randomly permuting the CES-D scores among participants 100,000 times and recalculating the Spearman correlation after each permutation. For each pathway, the permutation P value was calculated with a plus-one correction as (b + 1)/(100,000 + 1), where b was the number of permutations yielding an absolute correlation coefficient at least as large as the observed value. P values were adjusted across all 389 pathways using the Benjamini–Hochberg procedure.

As a prevalence sensitivity analysis, the Benjamini–Hochberg adjustment was repeated after restricting the analysis to the 354 pathways with nonzero predicted abundance in at least 5 of the 17 participants. Unadjusted P values below 0.05 were treated as nominal exploratory signals, whereas pathway-level statistical significance was defined as a Benjamini–Hochberg-adjusted q value below 0.05.

**Supplementary Figure Legends**

**Fig. S1** Comparison of gut bacterial alpha diversity between the control and higher depressive symptom groups.

Alpha diversity was calculated after rarefaction to 26,666 reads per sample using observed features, Chao1 richness, Shannon diversity, Pielou’s evenness, and Faith’s phylogenetic diversity. Participants with a Center for Epidemiologic Studies Depression Scale (CES-D) score below 16 were assigned to the control group (salmon, n = 13), whereas those with a score of 16 or higher were assigned to the higher depressive symptom group (blue, n = 4). Each dot represents one participant. Boxes indicate the interquartile range, horizontal lines indicate the median, and whiskers indicate the minimum and maximum values. Unadjusted *p* values are shown above the corresponding comparisons. No statistically significant between-group difference was detected for any of the five indices.

**Fig. S2** Principal coordinate analysis of weighted UniFrac distances between the control and higher depressive symptom groups.

Principal coordinate analysis (PCoA) of weighted UniFrac distances was used to visualize differences in the abundance-weighted phylogenetic composition of the gut bacterial community. Participants with a Center for Epidemiologic Studies Depression Scale (CES-D) score below 16 were assigned to the control group (red, n = 13), whereas those with a score of 16 or higher were assigned to the higher depressive symptom group (blue, n = 4). Each point represents one participant. Axis 1, Axis 2, and Axis 3 explained 43.32%, 17.73%, and 10.92% of the variation, respectively. Permutational multivariate analysis of variance detected a significant difference in community composition between the groups (pseudo-*F* = 2.42, *p* = 0.040, *q* = 0.040).

**Fig. S3** LEfSe cladogram of bacterial lineages associated with the control and higher depressive symptom groups.

The cladogram shows the taxonomic distribution of bacterial lineages identified by linear discriminant analysis effect size (LEfSe). Participants with a Center for Epidemiologic Studies Depression Scale (CES-D) score below 16 were assigned to the control group (red, n = 13), whereas those with a score of 16 or higher were assigned to the higher depressive symptom group (green, n = 4). From the center outward, the concentric rings represent the phylum, class, order, family, genus, and species levels. Node size reflects relative abundance. Red and green nodes and branches indicate taxa enriched in the control and higher depressive symptom groups, respectively, whereas yellow nodes indicate taxa that were not identified as differentially represented by LEfSe. The labeled lineages are (a) Micrococcaceae, (b) Micrococcales, (c) Porphyromonadaceae, and (d) Enterococcaceae.

**Fig. S4** Principal coordinate analysis of PICRUSt2-inferred microbial functional profiles in the control and higher depressive symptom groups.

Principal coordinate analysis (PCoA) based on Bray-Curtis dissimilarities was used to visualize predicted microbial functional profiles generated using Phylogenetic Investigation of Communities by Reconstruction of Unobserved States 2 (PICRUSt2). The profiles were summarized according to Clusters of Orthologous Groups (COG), Enzyme Commission (EC) numbers, Kyoto Encyclopedia of Genes and Genomes Orthology (KO), and MetaCyc pathways. Participants with a Center for Epidemiologic Studies Depression Scale score below 16 were assigned to the control group (red, n = 13), whereas those with a score of 16 or higher were assigned to the higher depressive symptom group (blue, n = 4). Each point represents one participant, and the percentage of variation explained by each axis is shown in parentheses. Permutational multivariate analysis of variance detected significant differences between the groups for COG (pseudo-*F* = 3.26, *R*² = 0.179, *p* = 0.0218), EC (pseudo-*F* = 3.05, *R*² = 0.169, *p* = 0.0244), KO (pseudo-*F* = 3.07, *R*² = 0.170, *p* = 0.0210), and MetaCyc pathway profiles (pseudo-*F* = 3.05, *R*² = 0.169, *p* = 0.0185). Permutational analysis of multivariate dispersions detected no significant difference in dispersion for the KO or EC profiles (*p* = 0.552 and *p* = 0.709, respectively).
